# Peptides derived of kunitz-type serine protease inhibitor as potential vaccine against experimental schistosomiasis

**DOI:** 10.1101/634360

**Authors:** Juan Hernández-Goenaga, Julio López-Abán, Anna V. Protasio, Belén Vicente Santiago, Esther del Olmo, Magnolia Vanegas, Pedro Fernández-Soto, Manuel Alfonso Patarroyo, Antonio Muro

## Abstract

Schistosomiasis is a significant public health problem in sub-Saharan Africa, China, South-East Asia and regions of South and central America affecting about 189 million people. Kunitz-type serine protease inhibitors have been identified as important players in the interaction of other flatworm parasites with their mammalian hosts. Here, we evaluate the protective efficacy of chemically synthesized of T- and B-cell peptide epitopes derived from a kunitz protein from *Schistosoma mansoni*. Putative kunitz-type protease inhibitor proteins were identified in the *S. mansoni* genome and their expression analyzed by RNA-seq. Gene expression analyses showed that the kunitz protein Smp_147730 (Syn. Smp_311670) was dramatically and significantly up-regulated in schistosomula and adult worms when compared to the invading cercariae. T- and B-cell epitopes were predicted using bioinformatics tools, chemically synthesized and formulated in the Adjuvant Adaptation (ADAD) vaccination system. BALB/c mice were vaccinated and challenged with *S. mansoni* cercariae. Kunitz peptides were highly protective in vaccinated BALB/c mice showing significant reductions in recovery of adult females (89-91%), and in the numbers of eggs trapped in the livers (77-81%) and guts (57-77%) of mice. Moreover, liver lesions were significantly reduced in vaccinated mice (64-65%) compared to infected control mice. The vaccination regime was well tolerated with both peptides. We propose the use of these peptides, alone or in combination, as reliable candidates for vaccination against schistosomiasis.

## Introduction

Human schistosomiasis is a water-borne debilitating disease caused by a trematode of the genus *Schistosoma*. It is estimated that 240 million people worldwide are infected with *Schistosoma* spp. which causes the loss of 1.5 million DALYs (Disability Adjusted Life Years) per year (1). In 1994, the WHO (World Health Organization) together with the *Schistosoma* Genome Network started a project aimed to sequencing the *S. mansoni* genome, which was published in 2009 (2) alongside the *Schistosoma japonicum* genome (3). Three years later the genome of *S. haematobium* was described (4). Schistosomes’ genome size is relatively large 409.5 Mbp for *S. mansoni*, 376 Mbp for *S. haematobium* and 403 Mbp for *S. japonicum* due to the presence of a large number of repetitive sequences (40-45%).

In recent years, high-throughput (next generation) sequencing technologies have provided a large amount of data on covering different aspects of schistosome biology. For example, genome sequencing of multiple isolates has revealed the complex population biology of schistosomes (5–6), and RNA-seq transcriptomic studies have allowed a better understanding of the gene expression patterns during these parasites’ life cycle (7–12). These data are made available to the research community via databases such as GeneDB, SchistoDB and WormbaseParasite (13–15).

The most interesting schistosome proteins are those related to host-parasite interactions (16), since they are accessible to the effector mechanisms of the host’s immune system and may be targets for development of drugs and vaccines against these helminths. There are two promising groups: parasite surface proteins and excretory-secretory proteins. The latter category includes several proteases (serine, cysteine and aspartic proteases) (17) as well as some protease inhibitors that ensure the survival of the parasite by inhibiting host proteases enzymes (18). MEROPS, a database of proteases and inhibitors, contains 1008 annotated entries for human proteases and homologs (19). The recent availability of the genome sequences of different mammals has allowed the identification of their entire protease composition, termed “degradome”, and its comparison with the human counterpart. The Degradome Database lists 569 human proteases and homologs classified into 68 families (20). A plethora of proteins has been proposed as potential vaccines against schistosomiasis, but only Sm14 and SmTSP-2 vaccines for *S. mansoni* have reached Phase I clinical trials and only the glutathione-S transferase rSh28GST (Bilhvax) against *S. haematobium* has reached Phase III (21).

Kunitz-type protease inhibitors belong to the family of serine protease inhibitors that are found in almost all organisms. They are small proteins containing around sixty amino acid residues (17) and have one or more kunitz motif: α + β with two β strands and two short α helices at the end of the domain. This domain also has three disulfide bonds between six conserved cysteines (22). Kunitz proteins have been involved in various physiological processes as blood coagulation, fibrinolysis, inflammation and ion channel blocking (17). However, there is limited information regarding kunitz-type protease inhibitors of parasitic helminths. These molecules have been described in *Fasciola hepatica* (23), *Echinococcus granulosus* (24) and *Ancylostoma* spp (25) and could to be reliable antigens for vaccine design. Kunitz-type protease inhibitors have been identified in the genomes of the three major *Schistosoma* spp, but only SjKI-1 from *S. japonicum* and SmKI-1 from *S. mansoni* have been expressed and functionally characterized (26–27). Recently, recombinant *S. mansoni* kunitz protein (rSmKI-1) formulated with Freund’s adjuvant was shown to induce partial protection against C57BL/b mice infected with *S. mansoni* (28). A strategy to design vaccines is based on the use of conserved peptides involved in critical physiological processes able to interact with major histocompatibility complex (MHC) class I and II molecules and drive protective immune responses. Minimal antigen epitopes with 13-18 amino acid long peptides can be designed to trigger B- and T cell immune responses and we can synthesize them chemically (29–30).

The aim of this study was to explore *S. mansoni* genome *in silico* to identify kunitz-type serine protease inhibitors and to study their expression profile in different life stages by RNA-seq, and to compare them with kunitz protein sequences from other schistosomes and other helminths. One kunitz T- and B-cell epitope were predicted, chemically synthesized and further tested as potential vaccine candidates against *S. mansoni* in mice

## Materials and Methods

### Animals and parasites

Seven-week-old SPF female BALB/c mice (Charles River, Lyon, France) weighing 18-20 g were allocated in standard cages with food and water *ad libitum*, light/dark cycle of 12/12 h and 22-25°C. Animal procedures complied with the Spanish (L 32/2007, L 6/2013 and RD 53/2013) and the European Union (Di 2010/63/CE) regulations. The Ethics Committee of the University of Salamanca approved animal use protocols (Ref. 15/0018). The size of groups was calculated by power analysis using “size.fdr” package in R and following the 3Rs recommendations (31–32). The animals’ health status was monitored during the experiments according to FELASA guidelines. *S. mansoni* was maintained in *Biomphalaria glabrata* snails as intermediate hosts and CD1 mice as definitive hosts. The number of cercariae and their viability were determined using a stereoscopic microscope (Olympus SZX9, Japan).

### Kunitz-type protease inhibitors study in *Schistosoma mansoni* genome

Amino acid sequences of all putative kunitz domain containing proteins of *S. mansoni* were retrieved from GeneDB and SchistoDB (14–15). Sequences containing at least six cysteines were kept for further analyses and aligned with other kunitz proteins from *S. japonicum*, *S. haematobium, E. granulosus*, *E. multilocularis* and *F. hepatica* available from GeneDB, SchistoDB, GenBank (33) and WormBase ParaSite (13). Amino acid identity between sequences was analyzed using alignments generated with Clustal Omega on line web-server (34) and then visually edited with BioEdit software *v*7.1.3 (35). Potential secretory signal peptides were predicted with SignalP 4.1 online tool (36) with a D-cutoff value of 0.45. Transmembrane helix regions in the sequences were predicted using TMHMM server v2.0 (37). GPI-anchored potential was estimated using fragAnchor (38).

### Kunitz gene expression in *S. mansoni* in cercariae, schistosomula and adults

RNA-seq data from Protasio et al (7) was used to investigate the expression profile of proteins with kunitz motifs. Fastq files corresponding to samples with accession numbers ERR022873, ERR022874, ERR022876-78 and ERR022880-81 were retrieved from www.ena.ac.uk; reads were mapped to the latest version of the *S. mansoni* genome WBPS12 (https://parasite.wormbase.org/Schistosoma_mansoni_prjea36577/) using HISAT2 v2.1.0 (39) with default parameters except for “–no-mixed –no-discordant”. Output SAM files were converted, sorted and indexed using SAMTOOLS v1.9 (40). Gene annotation as GFF was obtained from Wormbase ParaSite and corresponds to the database release WBPS12. A GTF version of the annotation was produced using GFFREAD from the CUFFLINKS suite v2.2.1 (41) with options “-F -T”. Counts per gene were computed using FEATURECOUNTS from the SUBREAD v1.6.3 package (42) with default parameters except for “‒primary ‒fraction -t exon -g gene_id”. Counts per gene were further processed using DESeq2 v1.16.1 (43) and visualization of gene expression changes were produced using Integrative Genomics Viewer (IGV) (44) and Tror GGPLOT2 (45) implemented in R (46).

A touchdown PCR (TD-PCR) was developed using the Smp_147730 sequence (Syn. Smp_311670). The reaction was optimised in 25 μL reaction mix containing: 2 μL of DNA extracted from *S. mansoni* adults, 13 μL H_2_O, 2.5 μL 10x reaction buffer, 2.5 μL MgCl_2_ (25 mM), 2.5 μL, dNTPs MIX (25 mM/dNTP), 1 μL (10 pmol) of each primer and 0.5 μL of Taq-polymerase 2,5 U (iNtRON Biotechnology, Inc). The program consisted of one cycle at 94°C 1 min, six cycles 94°C for 20 sec and a touchdown program of 15 cycles with successive annealing temperature decrement from 65°C to 60°C for 30 sec with 1°C decrement with a final extension at 72°C 10 min performed in a Mastercycler Gradient (Eppendorf). The products were monitored using 1.5 % agarose gel electrophoresis stained with ethidium bromide, visualized under UV light and then photographed (Gel documentation system, UVItec, UK). The DNA insert obtained was sequenced by the Sanger method at the Sequencing Service of the University of Salamanca.

### B and T-cell peptide prediction and chemical synthesis from S. mansoni kunitz protease inhibitor

The genetic sequence of the proposed kunitz protease inhibitor gene Smp_147730 (currently Smp_311670 in WBPS12 - https://parasite.wormbase.org/Schistosoma_mansoni_prjea36577/) was analysed *in silico* to identify potential B-and T-cell epitopes that could be soluble and easy to manufacture. Peptide SmKT was designed to induce a good T-cell response using the SYFPEITHI database (47) and the Immune Epitope Database (IEDB) (48), good MHC class II binders were searched for murine H2-E^d^ and human HLA-DRB1. Sequences with scores more than 20 in predictions based on k-mers of length 15 were selected. The BebiPred server, based on hidden Markov models (HMM), was used for predicting linear B-cell epitopes (BebiPred 1.0b). Prediction score is based on hydrophilicity and secondary structure prediction (49). The Predicted linear B-cell epitope was compared with the results found using the ANTHEPROT 3D software, which takes antigenicity, hydrophobicity, flexibility and solvent accessibility into account (50). A 20 amino acids region displaying the best score for each protein was selected as promising linear B-cell epitopes.

The predicted T and B-cell epitopes (referred to as SmKT and SmKB respectively) were chemically synthesized at Fundación Instituto de Inmunología de Colombia (FIDIC) (Bogotá, Colombia) by the solid-phase peptide synthesis according to Merrifield (51) and Houghten (52) using the t-Boc strategy and α-benzyhydrylamine (BHA) resin (0.7 meq/mg). One cysteine and a glycine residue were added at both amino and carboxyl-terminal ends to allow their polymerization via oxidization. Peptides were purified by reverse phase high performance liquid chromatography characterized by MALDI-TOF mass spectrometry and lyophilized. Freeze-dried synthetic peptides were re-suspended in phosphate buffered solution (PBS) and concentrations were determined with a BCA kit of Pierce (Rockford, IL). Peptide toxicity was determined in J774.2 mouse peritoneal macrophage cell line cultures. Cell viability was measured by CytoTox 96® Non-Radioactive Cytotoxicity Assay (Promega) (53).

### Vaccination trial using the ADAD vaccination system

Twenty-one female BALB/c mice were randomly allocated in four groups: Untreated and uninfected group (n=3), Adjuvant treated and infected group (AA0029+Qs) (n=6), SmKT vaccinated and infected group (AA0029+Qs+SmKT) (n=6) and SmKB vaccinated and infected group (AA0029+Qs+SmKB) (n=6). Mice received three vaccinations at 2-week intervals. SmKT and SmKB were formulated in the Adjuvant Adaptation (ADAD) vaccination system with non-hemolytic saponins from *Quillaja saponaria* (Qs; Sigma) and the synthetic aliphatic diamine AA0029 emulsified in a non-mineral oil (Montanide ISA763A, SEPPIC) with a 70/30 oil/water ratio. The ADAD vaccination system is administered using two subcutaneous injections. The first injection containing AA0029 and Qs emulsified in Montanide and the second injection, administered 5 days after the first, contains the antigen with AA0029 and Qs in the emulsion oil. Individual doses per injection in mice included 100 μg of AA0029, 20 μg of Qs and 10 μg of either SmKT or SmKB in a final 100 μL volume of emulsion with Montanide (54–55). Mice were weighed and monitored for signs of anaphylactic shock, erythema at the injection point and changes in behavior. Vaccinated and infection control mice were percutaneously challenged with 150±8 *S. mansoni* cercariae two weeks after the third vaccination. Mice were restrained with a mixture of 50 mg/kg ketamine (Imalgene1000, Merial), 5 mg/kg diazepam (Valium10, Roche Farma SA) and 1 mg/kg atropine (B. Braun, Madrid) administrated intraperitoneally. The abdomen was shaved and wetted with sterile water and then exposed to cercariae for 45 minutes using a ring (55). All mice were euthanized with a lethal dose of 60 mg/kg of pentobarbital plus 2 IU/mL of heparin and then perfused aseptically with PBS and heparin (500 IU/L). Paired worms as well as single males or females were obtained from the portal and mesenteric veins by portal perfusion at 8 weeks post-challenge. The liver and small intestine were digested in 5% KOH (w/v) overnight at 37°C with shaking and eggs per gram were counted three times using a McMaster chamber by two different researchers. The spleen, gut and liver weights were recorded. Liver injury was assessed by the number of granulomas in the surface determined by two pathologist independently using three micrographs (Olympus SZX9) and ImageJ 1.45 software (56).

### Humoral immune response by ELISA

Soluble *S. mansoni* adult worm antigens (SoSmAWA) were prepared for ELISA (55). Twenty adult worms per mL were suspended in sterile PBS with a protease inhibitor cocktail (Complete Mini EDTA-Free, Roche 04 693 159 001). The mixture was homogenized, frozen and thawed, sonicated and then centrifuged at 30,000 *g* for 30 min at 4°C. Supernatant protein concentration was determined using Micro BCA Protein Assay Kit. Blood samples were collected from mice before immunization and infection, and at the necropsy and analysed by indirect ELISA to detect specific IgG, IgG1 and IgG2a antibodies anti-SmKT, -SmKB and -SoSmAWA. A Corning Costar 96-well microplate (Cambridge, MA) was coated with 1 μg/mL of each peptide and SoSmAWA. The plates were then blocked with 2% of bovine serum albumin (Sigma) in PBS with 0.05% Tween 20 (PBST) for 1 h at 37°. Sera samples diluted at 1:100 in PBST were added in duplicate wells and incubated 1 h 37°C. Goat anti-mouse IgG-HRP, IgG1-HRP or IgG2a-HRP conjugates (Sigma) were used at 1:1000 in PBST and incubated 1 h at 37°C. The plates were washed and developed adding H_2_O_2_ (0.012%) and orthophenylenediamine substrate (0.04%) in 0.1 M citrate/phosphate buffer pH 5.0. The reaction was stopped with 3 N H_2_SO_4_ and read at 492 nm on a MultiSkan GO ELISA plate reader (Thermo Fisher Scientific, Vantaa).

### Parasitological and immunological data analyses

Data were expressed as the mean and standard error of the mean (SEM) and were tested for normality by the Kolmogorov-Smirnov test and homogeneity of variance by the Bartlett test. A one-way ANOVA test and multiple *post-hoc* comparisons with Tukey’s honest significance tests (HSD) or Kruskal-Wallis (K-W) tests were performed to analyze statistical differences among groups. A value of *P*<0.05 was considered statistically significant. Statistical analyses were performed with SIMFIT Statistical Package 7.4.1 (Manchester University, U. K. https://simfit.org.uk) and SPSS 21 software (IBM).

## Results

While the initial identification of kunitz domain containing proteins in *S. mansoni* was performed using the former v5.0 of the *S. mansoni* genome assembly (7), an updated and improved version the assembly was released during the production of this manuscript. Gene accession numbers have change between these two versions. The original accession numbers (found in v5.0) used to access nucleotide and amino acid sequences in different steps described in the methods of this manuscript were maintained as much as possible to allow cross-referencing with existing literature. However, wherever possible and appropriate, bioinformatics analyses were updated to confirm previous results against the new genome assembly and annotation (WBPS12, https://parasite.wormbase.org/Schistosoma_mansoni_prjea36577/).

### Kunitz-type protease inhibitors study of *Schistosoma mansoni*

The *S. mansoni* genome and gene annotation repositories (GeneDB and SchistoDB) were searched for all kunitz protein sequences. A total of 11 sequences were retrieved with putative kunitz domains in the *S. mansoni* genome. Only three sequences of potential interest remain in V 5.0: Smp_147730, Smp_139840 and Smp_012230. After a close analysis of their amino acid sequence only Smp_147730 (currently Smp_311670.1) contained a bona fide kuntiz-domain identified between residues 26 to 79 with the six highly conserved cysteine residues capable to establish three disulphide bonds, found in the range of 50-70 amino acids. In addition, a signal peptide was predicted in the first 21 amino acids of Smp_147730 (Syn. Smp_311670) with a D-score of 0.855 according with SignalP 4.1 (36). The cleavage site was located between positions 20-21 where Y-score showed the highest value (Y=0.828). In consequence, residues 1-20 were removed from the B- and T-cell peptide prediction. No transmembrane or GPI anchor domains were found with TMHMM server v2.0 (57).

A new version of the *S. mansoni* genome (version 7, unpublished, available from https://parasite.wormbase.org/Schistosoma_mansoni_prjea36577/Info/Index/, database version WBPS12) was released during the preparation of this manuscript. The gene Smp_147730 has been renamed Smp_311670 and it is predicted to produce two alternative transcripts. The sequence of Smp_311670.1 is identical to our confirmed kunitz protein sequence while Smp_311670.2 represents a longer alternative transcript.

### Comparison of Smp_147730 (Syn. Smp_311670) with trematode kunitz proteins

The Smp_147730 (Syn. Smp_311670) sequence was compared to other putative kunitz proteins of Platyhelminthes identified in sequence databases. There were seven sequences retrieved from GeneDB of *S. japonicum* but only four had a six-cysteine kunitz domain with an identity ranging between 26.05 and 42.03% (Fig. 1A). Two out of eight *S. haematobium* sequences present in SchistoDB did not include a kunitz domain and the identity of the remaining proteins to Smp_147730 (Syn. Smp_311670) ranged between 18.05 and 74.62% (Fig. 1B). Seven kunitz protein sequences in GeneBank were attributed to *E. granulosus* but only five have a kunitz domain and identities ranged 15.45 to 37.33% (Fig. 1C). There were six sequences sharing the domain available in GeneBank from *E. multilocularis* with identities from 14.41 to 35.80% out a total of eight retrieved (Fig. 1D). Three identical *F. hepatica* sequences were identified in WormBase ParaSite each presenting a kunitz domain and sharing 32.93% residue identity with Smp_147730 (Fig. 1E).

**Figure 1.**
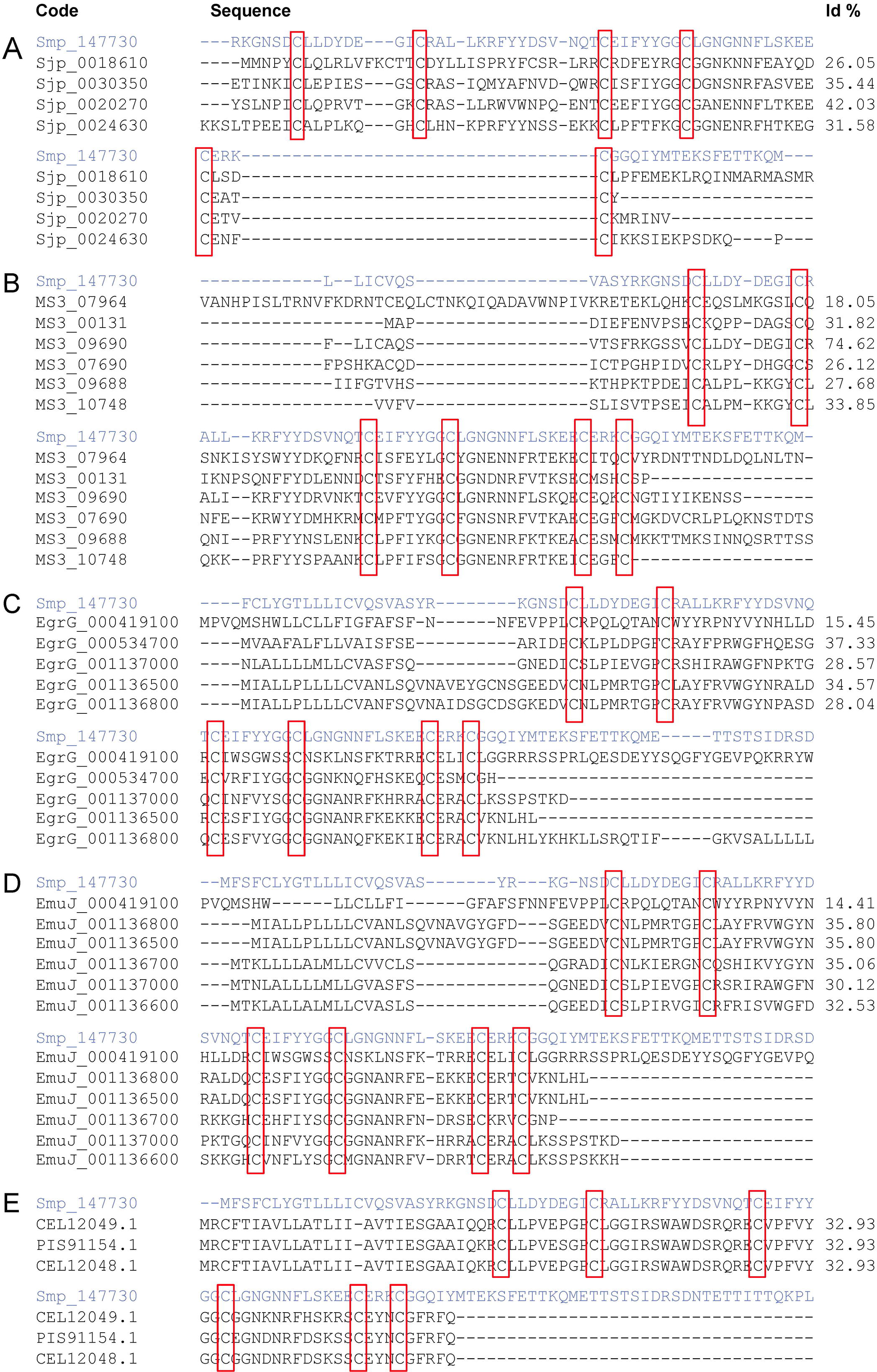
Comparison of sequence Smp_147730 (Syn. Smp_311670) from *S. mansoni* in a multiple sequence alignment with kunitz-type proteins from *S. japonicum* (**A**), *S. haematobium* (**B**), *Echinoccocus granulosus* (**C**), *E. multilocularis* (**D**) and *Fasciola hepatica* (**E**). Kunitz domain positions are highlighted in red boxes.

### Transcriptome analysis and differential expression of Smp_147730 (Syn. Smp_311670) kunitz gene

Smp_311670 is located in Chromosome 2 (37,805,700 and 37,811,500, forward strand) of the WBPS12 *S.mansoni* genome assembly. Transcriptome analysis by RNA-seq showed that Smp_311670.2 was significantly up-regulated (adjusted p-value < 0.01) in 24 hours schistosomula and adult worms with respect to cercariae. (Fig. 2, Supplementary Table 1. No significant difference was found between cercariae and 3-hours-schistosomula (Supplementary Table1).

**Figure 2.**
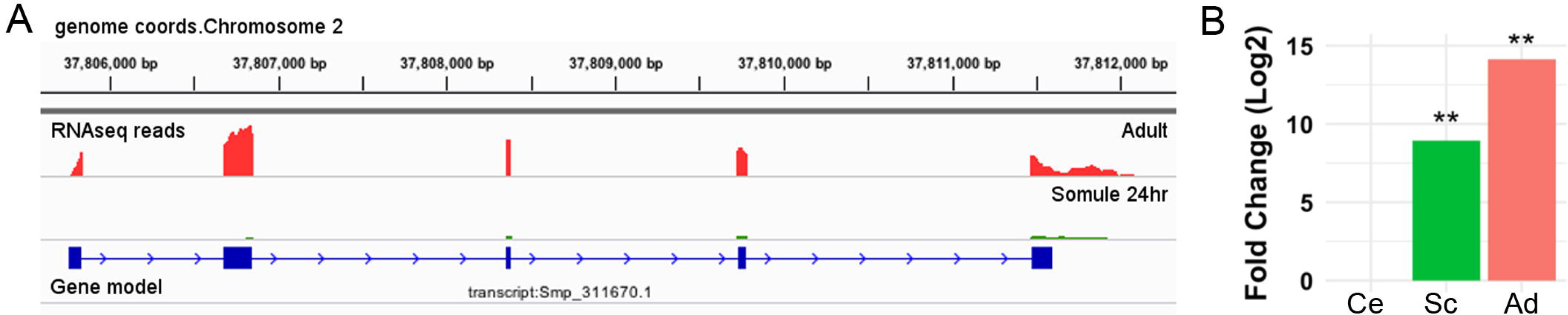
Visualisation of Smp_311670.1 in its genome context and gene expression. (**A**) Coverage plots for adult worms and 24-hours-schistosomula are shown using Integrative Genomics Viewer (IGV). Three-hours schistosomula and cercariae are omitted for clarity. (**B**) Relative gene expression of Smp_311660.2 is represented as a Log2 Fold Change relative to cercariae (Ce), 24-hour-schistosomula (Sc) and adult (Ad), ** *P* < 0.01.

Primers were designed to amplify Smp_147730 DNA (Syn. Smp_311670) sequence using *S. mansoni* adult DNA, Forw. 5’-TACTGACAGGGCTCACTACGCT-3’ and Rev. 5’-ACGCTCGCCTTCACACCCC-3’ by TD-PCR, the amplified region spanned exon 1 and 2. A 1444 bp insert was obtained and purified by agarose gel electrophoresis, quantified and sequenced (Fig. 3A). A consensus sequence was obtained using Bioedit software and compared through BLAST to sequences recorded in databases.

**Figure 3.**
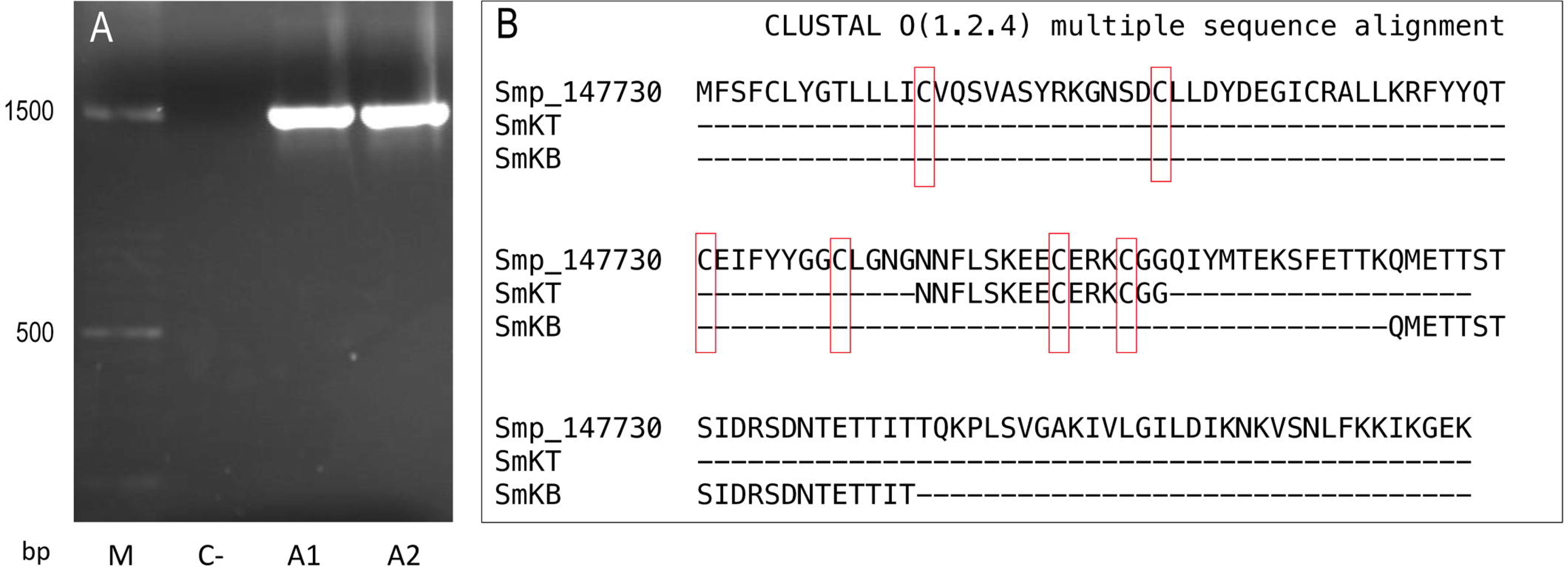
Sequences of Smp-147730 (Syn. Smp_311670) of *Schistosoma mansoni*:(**A**) Agarose gel electrophoresis insert of 1444 base pairs (bp) obtained by PCR from two adult DNA samples (A1 and A2) with negative control (C-) and mass scale (mM). (**B**) Amino acid sequence and alignment with Clustal Omega of predicted T-cell peptide (SmKT) and B-cell peptide (SmKB).

### T- and B-cell epitope prediction and toxicity assessment

The online servers SYFPEITHI (http://www.syfpeithi.de (47)) and Immune Epitope Database (IEDB) (http://www.immuneepitope.org/ (48)) resulted in the prediction of a 15-amino acids T-cell peptide (SmKT) in position 66-80 of Smp_147730 (Syn. Smp_311670) with a score of 22 for H2-Ed and score of 20 for HLA-DRB1*0401 inside the kunitz domain (Fig 3B). A 20-amino acid B-cell peptide (SmKB) was predicted located on residues 94-114 by BepiPred server (http://www.cbs.dtu.dk/services/BepiPred (47)) and ANTHEPROT (http://antheprot-pbil.ibcp.fr (50)) outside of the kunitz domain (Fig 3B). Peptides were obtained with purity more than 90%. Each peptide was assayed ranging from 1 to 50 μg/mL for *in vitro* cytotoxicity evaluation to J774.2 mouse macrophages. Results showed that more than 90% of macrophages were still viable after three days of treatment with SmKT and SmKB peptides in all conditions.

### Vaccination with SmKT and SmKB in ADAD with AA0029 triggers protection against *S. mansoni* infection

The capacity of SmKT and SmKB to induce protection in BALB/c mice against *S. mansoni* infection was evaluated. SmKT, T-cell peptide, formulated in ADAD with the synthetic immunomodulator AA0029 induced higher levels of protection measured by worm recovery with especially high reduction in the number of female worms collected by perfusion (91%; *P* = 0.0002) (Table 1). Also, significant reduction in number of eggs present in the liver (77%; *P* = 0.0044) and in the small intestine (57%; *P* = 0.0208) were detected (Table 1). Liver damage evaluated by the numbers of granulomas on the hepatic surface was also significantly reduced (65%; *P* = 0.0041) compared with controls (Figure 4). BALB/c mice immunized with the SmKB B-cell peptide, also showed a high reduction in female worms (89%; *P* = 0.0003) (Table 1), pronounced decreases in the number of eggs present in the liver (81%; *P* = 0.0030) and in the intestine (77%; *P* = 0.0028) (Table 1), as well as a reduced number of granulomas in the liver (64%; *P* = 0.0044) (Figure 4). Both vaccine candidates (SmKT and SmKB) showed comparable protection in terms of eggs trapped in tissues and liver lesions. No signs of anaphylactic shock, erythema or changes in behavior were observed in vaccinated mice with either peptide. No mice died during the trial. Slight subcutaneous traces of the emulsion were observed at the point of injection.

**Figure 4.**
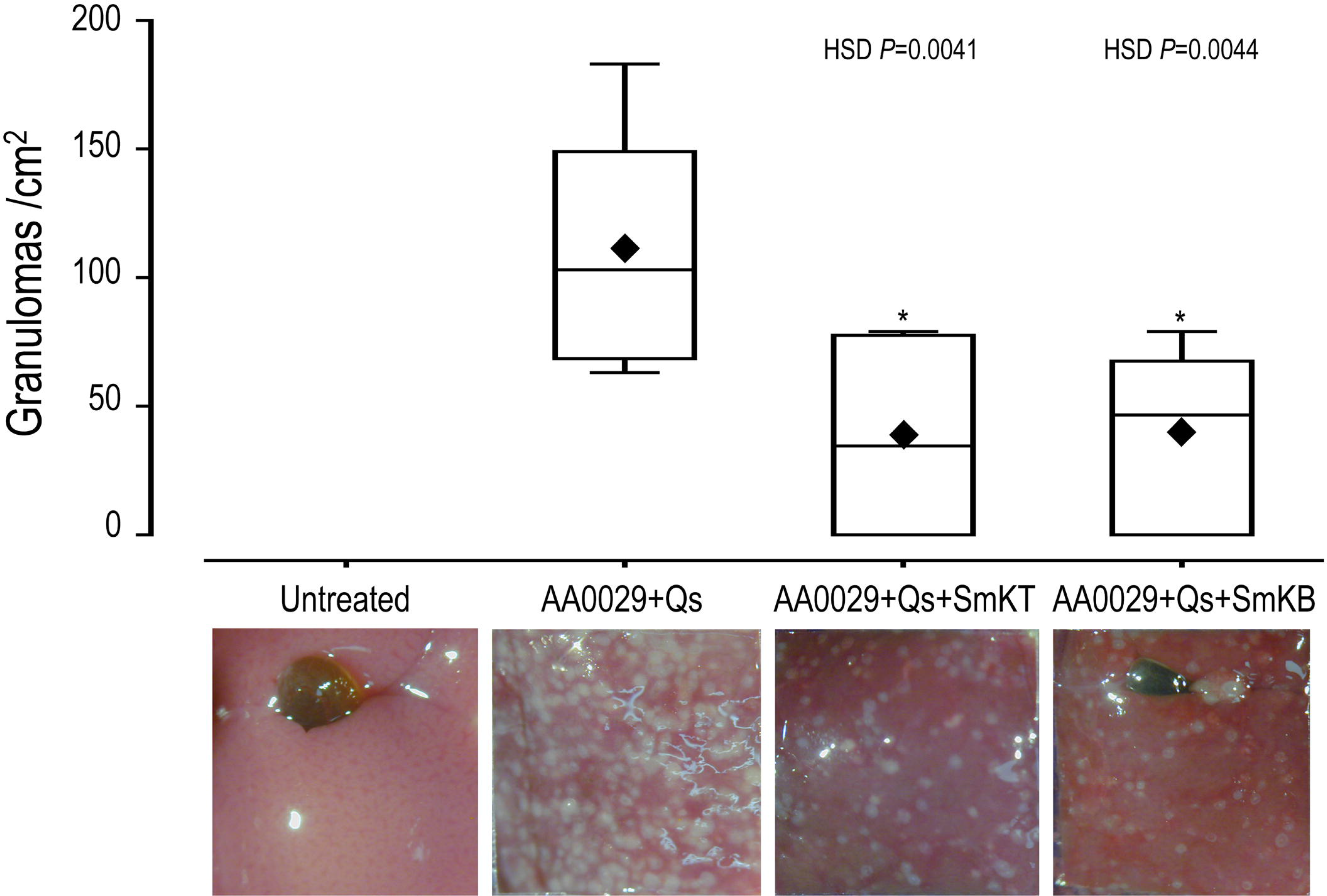
Effect on liver lesion of vaccination with SmKT and SmKB formulated in the Adjuvant Adaptation (ADAD) vaccination system with the synthetic immunomodulator AA0029 and *Quillaja saponaria* saponins (Qs) in BALB/C mice challenged with 150 cercariae of *S. mansoni*. ANOVA F_(3,17)_ 7.246 and p>0.0024, and post-hoc Tukey’s honest significance different (HSD) test *P* values are depicted in the chart. A representative micrograph of each group was included. ♦ represents means.

**Table 1.** Effect of vaccination with SmKT and SmKB formulated in the adjuvant adaptation (ADAD) vaccination system on total female and male worms counts, and eggs per gram (EPG) trapped in the tissues of BALB/C mice challenged with 150 cercaria of *S. mansoni*. Percentage of reduction (R). Data are presented as the mean and standard error of the mean (SEM). ANOVA and post-hoc Tukey’s honest significance test were used.

| Groups | Female worms<br>(mean ± SEM) | R<br>(%) | Male worms<br>(mean ± SEM) | R<br>(%) | EPG in liver<br>(mean ± SEM) | R<br>(%) | EPG in gut<br>(mean ± SEM) | R<br>(%) |
| --- | --- | --- | --- | --- | --- | --- | --- | --- |
| AA0029+Qs | 11.5 ± 2.8 | - | 7.7 ± 1.8 | - | 11050 ± 2928 | - | 11223 ± 1609 | - |
| AA0029+Qs+SmKT | 1.0 ± 0.4* | 91 | 10.7 ± 3.4 | NR | 2561 ± 1510* | 77 | 4872 ± 2666* | 57 |
|  | *P*= 0.0002 |  |  |  | *P*= 0.0044 |  | *P*= 0.0208 |  |
| AA0029+Qs+SmKB | 1.3 ± 0.4* | 89 | 21.7 ± 4.0 | NR | 2113 ± 700* | 81 | 2529 ± 933* | 77 |
|  | *P*= 0.0003 |  |  |  | *P*= 0.003 |  | *P*= 0.0028 |  |
| ANOVA | *F*<sub>(3,17)</sub>= 10.721 |  | *F*<sub>(3,17)</sub>=6.787 |  | *F*<sub>(3,17)</sub>= 6.243 |  | *F*<sub>(3,17)</sub>= 6.112 |  |
|  | *P*= 0.0003 |  | *P*= 0.0033 |  | *P*= 0.0047 |  | *P*= 0.0052 |  |

\* Significant differences in comparison with AA0029+Qs controls. NR no reduction

### Immunogenicity of *S. mansoni* kunitz SmKT and SmKB peptides and immune response against SoSmAWA by ELISA

Indirect ELISA tests were performed to examine the ability of SmKT and SmKB to induce humoral immune responses. A significantly higher production of specific anti-SmKB IgG was observed in the AA0029+Qs+SmKB vaccinated group compared to uninfected group after the second immunization (*P* = 0.0030), which was maintained until the end of the experiment (Figure 5A). By contrast, no increase of specific anti-SmKT IgG antibodies was observed in AA0029+Qs+SmKT or in AA0029+Qs+SmKB vaccinated mice (Figure 5B). Although antibodies to SmKT were detected after challenged, these were lower than those elicited by SmKB (Figure 5).

**Figure 5.**
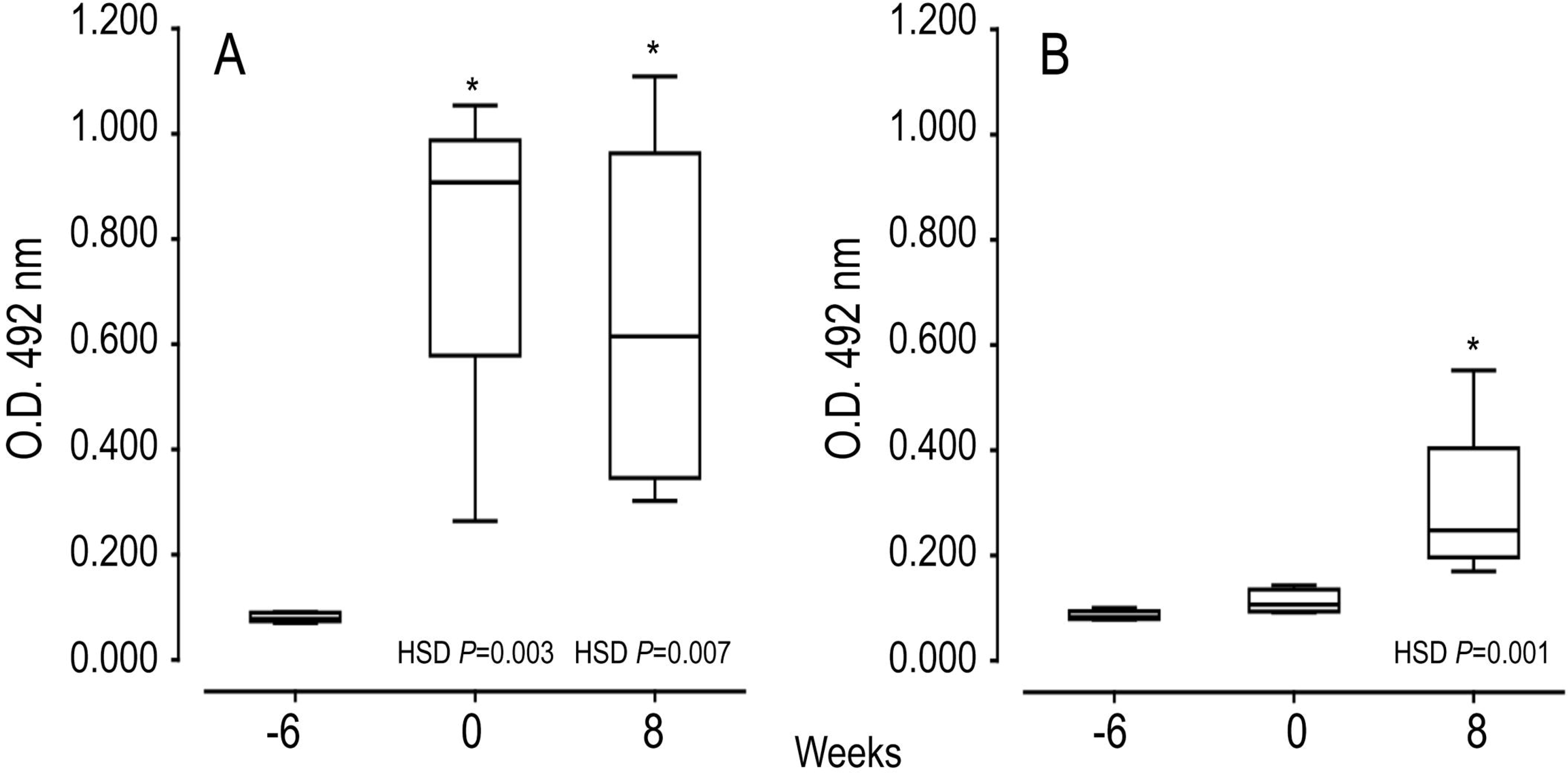
Serum antibody responses by ELISA during vaccination trials to SmKB (**A**) ANOVA F_(2,15)_ 10.910 *P* = 0.0017) and SmKT (B) ANOVA F_(2,15)_ 10.672 *P* = 0.0013) of mice vaccinated with AA0029+Qs+SmKB and AA0029+Qs+SmKT respectively. BABL/c mice were vaccinated using the adjuvant adaptation (ADAD) vaccination system and then challenged with 150 cercariae of *S. mansoni*. Data are presented as mean and standard error of the mean. * p<0.05 compared to serum sample before treatments.

Specific IgG, IgG1 and IgG2a responses against soluble worm antigen (SoSmAWA) were studied using an indirect ELISA. All infected groups showed an increase in total IgG production to SoSmAWA at 8 weeks post-challenge (0.584±0.118–0.720±0.107) compared to the uninfected group (0.250±0.064). All infected groups showed a significant higher production of IgG1 (ANOVA F_(3,17)_ 4.565 *P*=0.0160) at 8 weeks post-challenge (0.746±0.117−0.851±0.107) in comparison with uninfected group (0.263±0.005). By contrast, no significant increase of IgG2a antibodies to SoSmAWA was found during the experiment compared with uninfected group.

## Discussion

Important progress combating schistosomiasis have been made from 2013 to 2016 as reflected in the reduction of case numbers from 290 to 190 million (1, 58). This decrease in disease burden was mainly achieved through mass preventive chemotherapy with large-scale praziquantel administration complemented with safe water supplies, sanitation, hygiene education and snail control. There is increasing pressure for the developing of new anti-schistosomiasis drugs. Praziquantel, the drug of choice for schistosomiasis, has limitations because it acts only against the adult stage of the schistosome life cycle. In addition, there are mayor concerns regarding the emergence of drug resistance and/or reduced susceptibility to praziquantel due to its extensive use (59). Meanwhile, the development of a vaccine against this parasite is still high in the research agenda because it would complement the use of praziquantel to reduce disease, stop transmission and eradicate the disease. Purified or recombinant proteins from schistosomes, host-parasite interface antigens in tegument or gastrodermis, or genome mining by reverse vaccinology have been tested as vaccine candidates but only glutathione-S transferase rSh28GST (Bilhvax) have reached Phase III clinical trials (60).

Schistosome kunitz-type serine protease inhibitors have been associated to successful invasion, migration and development of the parasite in their host. They act by neutralizing the destructive action of host proteases on the invading schistosome (61). In other trematodes the secretion/release of proteins with kunitz domains interfere with the maturation of host dendritic cells and regulate host proteases resulting in impairment of defense responses (17). The kunitz protein SmKI-1 isolated from *S. mansoni* was found in excretory-secretory products and tegument of adults as well as eggs. It was observed that it also impairs neutrophil chemotaxis and elastase activity, coagulation and inflammation mechanisms in the host inducing immune evasion to ensure their survival. Moreover, the recombinant SmKI-1 delayed blood clot formation, inhibited several trypsin proteases but had no effect on pancreatic elastase or cathepsins (27). Therefore kunitz proteins are desirable new targets for vaccine development against schistosomes. Recombinant rSmKI-1 has previously been tested as a vaccine candidate as well as fragments involving the kunitz domain and the C-terminal tail (28, 62).

The potential of kunitz domain containing proteins as vaccines led us to study the published sequences of these genes in the three main schistosome species (63). We found 11 candidate DNA sequences containing the kunitz domain in several genome annotations of *S. mansoni* but only Smp_147730 (Syn. Smp_311670) had a six-cysteine residue characteristic of a *bona fide* kunitz domain. Also, we compared Smp_147730 (Syn. Smp_311670) with predicted kunitz type proteins available in database from *S. haematobium*, *S. japonicum*, *E. granulosus*, *E. multilocularis* and *F. hepatica*. The identity of the sequence with *S. haematobium* was the highest, up to 74% but in the other parasites was much lower, up 43%. This indicates that although the structure of kunitz domain was preserved in the different species these proteins could evolve separately and could be species-specific. These sequence differences correspond with the wide functional diversity of kunitz proteins in several species (64).

We focused on Smp_147730 (Syn. Smp_311670), studying its expression by RNA-seq and its identification by PCR in the *S. mansoni* strain maintained in our laboratory. We observed high expression of Smp_147730 (Syn. Smp_311670) after the transformation from cercaria to schistosomulum and even higher expression in the adult stage suggesting a role in schisosomulum development and the prolonged exposure in portal mesenteric veins of adults. The skin- or lung- migrating schistosomula and adult stages are regarded as major targets to design vaccines against schistosomes (65). With this in mind, our strategy was to design new synthetic high affinity peptide candidates composed of a short chain of amino acids containing the specific antigen determinant against functional regions that the parasite needs to survive (66). Several epitopes included in a vaccine would trigger humoral and cellular protective response using an adequate adjuvant or delivery system (67). We designed a T-cell peptide of 15 amino acids (SmKT), candidate from Smp_147730 sequence (Syn. Smp_311670), able to stimulate mouse and human MHC class II and a linear B-cell peptide of 20 amino acids (SmKB) based in physicochemical properties able to produce a humoral response. These *in silico* analyses are considered feasible, fast, and accurate in designing subunit vaccines against infectious diseases and could produce safer vaccines that are easier to manufacture and store than conventional ones (68). We formulated these two candidate peptides in the Adjuvant Adaptation (ADAD) vaccination system because adjuvants are recognized to have crucial importance in vaccine development. This adjuvant approach is a long-term delivery system feasible to use in vaccine development against helminths overcoming the issues of the experimental Freund’s adjuvant (54–55).

We next examined whether T- and B-cell epitopes could induce protection in BALB/c mouse experimental schistosomiasis. Both SmKT and SmKB candidates conferred a protection in terms of reduction in female worms, eggs trapped in tissues and liver lesions. These peptides could be useful to reduce liver granuloma pathology, and severe colonic damage and polyps. Fewer eggs in intestines could lead to less passage of eggs in feces and consequently could reduce transmission. The protection is higher than those obtained with the approaches of Morais et al (28) (34-43%) and Ranasinghe et al (69) (36-47%) using the whole recombinant rSmKI-1 or with the C-terminal-tail fragment (28-30%) (28) using Quil A with CBA mouse model or Freund’s adjuvant and C57BL/6 mice. While these different levels of protection could be explained by differences in adjuvant and animal model they all indicate the potential of Smp_147730 (Syn. Smp_311670) as good vaccine candidate. Our peptides seem to act against female worms leading to lower production of eggs and fewer lesions. Curiously our SmKT peptide of 15 -mer including only two cysteines of the conserved kunitz domain induced protection when the KI-fragment of 62 -mer involving three cysteines conserving the inhibitory activity against trypsin and neutrophil elastase tested by Morais et al (28) did not. Moreover, the SmKT induced weak antibody response with only significant increase at week 8 post-infection. This apparent incongruence could be related with the fact that conserved regions involved in critical biological functions for the parasite are poorly antigenic but interesting to develop immune response or to increase vaccine efficacy (30). On the other hand, SmKB peptide of 20 -mer and the C-terminal-tail fragment with 67 -mer containing the antiprotease activity used by Morais et al (28) indicating the value of B-cell mediated antibody response in schistosomiasis (62). Vaccination with SmKB induced a high production of specific IgG, contributing to control of adult phase as it was described in natural resistance to infection in people living in hyperendemic areas (70–72) and experimental models (73).

## Conclusion

Here we provide evidence for the protective capacity of two peptides SmKT and SmKB derived from kunitz proteins of *S. mansoni*. These peptides induced reduction in female worms, eggs in tissues and hepatic damage when administered subcutaneously formulated in the ADAD vaccination system. A single epitope vaccine could be insufficient to trigger a high level of protection, thus the combination with other synergic candidates in a multi-antigen vaccine must be tested in order to improve protection against *S. mansoni*.

## Acknowledgements

We thank Parasite Genomics Group from Welcome Sanger Institute, particularly to Matt Berriman (Group Leader) and James Cotton for the critical review of the draft. We thank the Sequencing Service and Animal Experimentation (Reg. # PAE/SA/001) facilities of the University of Salamanca. We thank Alba Torres Valle for her technical help in *S. mansoni* life cycle, ELISA and animal management. Montanide ISA763A was a gift of MSD Animal Health, Carbajosa de la Sagrada, Salamanca, Spain.

## Data Availability

Statement: All relevant data are within the manuscript and its Supporting Information files.

